# Defining phylogenetic relationships: a complete and minimal set of combinatorial properties for triplets

**DOI:** 10.1101/2024.03.19.585714

**Authors:** Rineau Valentin, Prin Stéphane

## Abstract

Triplets, as minimal informative rooted trees, are fundamental units of information in phylogenetics. Their importance for phylogenetic reconstruction, cladistic biogeography, or supertree methods relies on the fact that any rooted tree can be decomposed into a set of triplets. In order to formalize the tree building from a consistent triplet set, several *k* -adic rules of inference, i.e., rules that allow us to deduce at least one new triplet from exactly *k* other ones, have been identified. However, it remains unclear whether it is possible to reduce all the possible *k* -adic rules to a finite set of basic properties. In order to solve this problem, we propose here to define triplets in terms of degree of equivalence relations. Given the axiomatic definition of the latter, we establish a list of the most basic properties for triplets. With such an approach, we finally prove that the closure of any coherent triplet set can be computed uniquely from these basic properties.

## 1 Introduction

Representing affinity, proximity, or kinship relationships between objects by graph-theoretic trees and infering such trees is a problem at the heart of highly diverse fields, such as evolutionary biology, but also in biogeography, cultural evolution or even linguistics. An important property of trees is that they can be broken down into subtrees in order to be analysed and recombined. In the field of evolutionary biology, the increase in biological data as well as in processing capacities has made it possible to generate more and more phylogenetic trees on various taxonomic groups. New problems have arisen with the increasing number of trees concerning the manipulation and amalgamation of several trees into a unified hypothesis in order to make their multitude intelligible, and, with them, new phylogenetic methods of tree reconstruction and combination (Bininda-Edmonds 2004). For example, supertree-like methods have become central for phylogeny estimation in the past ten years. As a result, a growing number of algorithms have been designed to solve supertree-like problems (Nguyen et al. 2012; Sevillya et al. 2016; Fleischauer and Bocker 2017). The development of these methods and practices implies more than ever to have a relevant formal and axiomatic description of trees and phylogenetic relationships as mathematical structures on which biologists can rely for tree manipulation and amalgamation.

Among the different types of phylogenetic trees, those that can be qualified as “cladistic” are of particular interest. A cladistic tree is any completely ramified and single-rooted tree such that only its leaves are labelled. In other words, a cladistic tree is a hierarchical tree expressing the degree of kinship relationship sensu Prin (2016) among the objects represented by its leaves. The degree of kinship relationship, also known as “cladistic relationship” (Camin and Sokal 1965; Hull 1970), “phylogenetic relationship” (Hennig 1965; Nelson 1994), “relationship of common ancestry” (Nelson 1970, 1972, 1973), or still “degree of relationship” (Platnick 1979), consists of saying two things: (i) a trivial (or non-informative) component, that is, given two or more distinct objects (e.g. taxa), any of them is always closely related to itself in respect to any other one distinct from it; and (ii) a non-trivial (or informative) component, that is, given at least three distinct items, there is a possibility that at least two of them are more closely related than either is to the other ones. For example, the cladistic tree (((*ab*)*cd*)*e*) trivially states that for all distinct *x, y* ∈ {*a, b, c, d, e*}, *x* is closely related to itself in respect to *y*, and non trivially states that *a, b, c*, and *d* are closely related in respect to *e*, and *a* and *b* are closely related in respect to *c, d*, and *e*. Conversely, the cladistic tree (*abcd*) only trivially states that for all distinct *x, y* ∈ {*a, b, c, d* }, *x* is closely related to itself in respect to *y*.

Hence, the smallest possible informative cladistic trees are those that state that given exactly three distinct objects, two of them are more closely related than either is to the third. For example, the cladistic tree ((*ab*)*c*) states that *a* and *b* are closely related in respect to *c*. This fact allows us to introduce triplets sensu Vach (1994) and Wilkinson (1994: 344) as the units of cladistic information. In phylogenetics, a triplet, also called a “triad” (Adams 1986: 302), a “three-item statement” (Nelson and Platnick 1991), or still a “rooted triple” (Bryant 2003: 3), is an abstract object establishing that given three disjoint taxa, two of them are more closely related than either is to the third (Nelson and Platnick 1991). More generally, any triplet consists of saying that given three distinct objects (of a given kind), two of them are equivalent compared to the third. For example, a possible triplet for the three distinct objects *a, b*, and *c* is the one asserting that *a* and *b* are equivalent in respect to *c*. If appropriate, one writes “(*ab*)*c*” or “*ab*|*c*”. Note also that the order between *a* and *b* is irrelevant, i.e. (*ab*)*c* = (*ba*)*c*. Any triplet is thus formally equivalent to a minimal informative cladistic tree and conversely. For example, ((*ab*)*c*) and (*ab*)*c* corresponds to each other.

It is always possible to deduce a single triplet set *T* (*G*) from any cladistic tree *G* as follows: for all leaves *a, b, c* of *G*, (*ab*)*c* is in *T* (*G*) if and only if there is at least one node into *G* leading to *a* and *b* but not to *c*. For example, the cladistic tree (((*ab*)*cd*)*e*) leads to the triplet set {(*ab*)*c*, (*ab*)*d*, (*ab*)*e*, (*ac*)*e*, (*ad*)*e*, (*bc*)*e*, (*bd*)*e*, (*cd*)*e*}. In the same way, (*abcd*) leads to the empty set, since it implies no triplet. Conversely, it is possible, under some constraints, to build a cladistic tree from a triplet set. On this point, the fact that any cladistic tree *G* leads to a single triplet set *T* (*G*) means not only that *T* (*G*) represents *G* but also that as *G* satisfies some properties, it follows the same thing for *T* (*G*). From this, one may think that only complete triplet sets, that is, triplet sets containing all the triplets deducible from a given cladistic tree, could represent cladistic trees. However, it appears that cladistic trees can also be represented by incomplete triplet sets. For example, (*ab*)*d*, (*bc*)*d* can be considered as representing ((*abc*)*d*). Yet this set is well incomplete as ((*abc*)*d*) implies the complete triplet set {(*ab*)*d*, (*bc*)*d*, (*ac*)*d* }. Such a thing is in fact possible only because triplets obey to several combinatorial properties. As a result, to determine whether there is a complete and minimal set of basic rules for combining triplets is of primary importance for tree amalgamation methods.

Historically, the combinatorial properties of triplets and quartets (minimally informative unrooted trees) have mostly been investigated as a problem of *k* -adic inference rules (e.g. Dobson 1974; Dekker 1986; Bryant Steel 1995; Wilkinson et al. 2004). Given a natural number *k* ≥ 2, a *k* -adic inference rule for triplets is a combinatorial property allowing us to deduce at least one new triplet from *k* previous ones. For example, (*ab*)*c* and (*bc*)*d* imply (*ab*)*d* and (*ac*)*d*. Thus, even if the cladistic tree (((*ab*)*c*)*d*) implies the complete triplet set {(*ab*)*c*, (*ab*)*d*, (*ac*)*d*, (*bc*)*d* }, it can be represented by the incomplete set {(*ab*)*c*, (*bc*)*d* }. In its investigation of such rules for quartets, Dekker (1986) discovered not only some dyadic rules but also several rules of higher order that cannot be reduced to a repeated application of the dyadic ones. From that, he conjectured the existence of an infinite set of such rules. Then, Bryant Steel (1995) proved this conjecture not only for quartets but also for triplets. More explicitly, they proved the following statement: given a natural number *k* ≥ 3, there is always at least one *k* -adic rule for combining triplets that cannot be deduced to rules of lower order. Hence the impossibility to reduce all the combinatorial properties of triplets to a finite set of *k* -adic rules.

The aim of this paper is to demonstrate that it is possible to reduce all the combinatorial properties of triplets to a finite set of basic properties. We do so by defining triplets in terms of degree of equivalence relations that are specific mathematical ternary relations axiomatically defined first by Colonius and Schulze (1981), and then by McMorris and Powers (2003). For this, we will first address triplets as the smallest possible non-trivial degree of equivalence relations in order to infer their most basic combinatorial properties; then, we will adress about the consistency of triplet sets and their ahograph-based representation; after that, we will address the concept of the closure of a consistent triplet set; finally, we will conclude on the possibility to compute such a closure only from the basic properties previously highlighted.

## 2 Triplets as degree of equivalence relations

### 2.1 Degree of equivalence relation

A triplet is an object stating that given three distinct items, two of them are more closely related to each other than any is to the third. This fact will allow us to formally define triplets in terms of mathematical (i.e., set-theoretic) ternary relation (on a given set). More precisely, we propose below to define a ny triplet as what we call a “degree of equivalence relation (on a given set)”. These relations were first axiomatically be defined by Colonius and Schulze (1981), and secondly by McMorris and Powers (2003) under the name of “hierarchical relation”. We propose here Prin’s (2016) axiomatic definition for our purpose.

Let *A* be a finite n on-empty s et. A (finite) *degree of equivalence relation* on *A* is any set *D* ⊆ *A*^3^ such that for all *w, x, y, z* ∈ *A* :

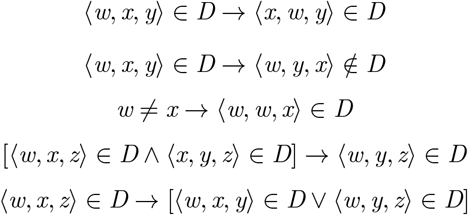

In other words, a degree of equivalence relation on a finite non-empty set is any ternary relation on that set which is respectively 1-2 symmetric, 2-3 asymmetric, 1-2 conditionally reflexive, 1-2 transitive, and 2-3 negatively transitive.

Now, let **D**(*A*) be the set of all degree of equivalence relations on *A*. For any *D* ∈ **D**(*A*) and any ⟨*x, y, z* ⟩ ∈ *D*, ⟨*x, y, z* ⟩ is *trivial* (or *non-informative*) if *x* = *y*, and *non-trivial* (or *informative*) otherwise. On this basis, the *trivial degree of equivalence relation* on *A* is the set *D* _0_(*A*) such that

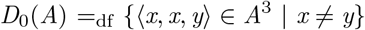

For any *D* ∈ **D**(*A*), it follows that

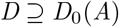

and

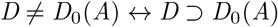

### 2.2 Formal definition and properties

We now give a formal definition of triplets as the smallest possible non-trivial degree of equivalence relations. Let *A* be a finite set such that |*A*| ≥ 3 and let three distinct *x, y, z* ∈ *A*. Then let

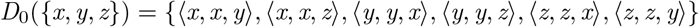

be the trivial degree of equivalence relation on {*x, y, z* }. In this case, one can define (*xy*)*z* as follows:

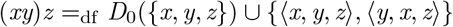

For all *u, v, w, x, y, z* ∈ *A* and any *D* ∈ **D**(*A*), it follows:

- (*uv*)*w* is deductible from *D* if and only if (*uv*)*w* ⊆ *D*
- (*uv*)*w* = (*xy*)*z* ↔ ({*u, v*} = {*x, y*} ∧ *w* = *z*), which implies (*uv*)*w* = (*vu*)*w*
- (*uv*)*w* ⊆ *D* → ¬[(*uw*)*v* ⊆ *D* ∨ (*wv*)*u* ⊆ *D*]
- [(*uv*)*x* ⊆ *D* ∧ (*vw*)*x* ⊆ *D*] → (*uw*)*x* ⊆ *D*
- (*uv*)*x* ⊆ *D* → [(*uv*)*w* ⊆ *D* ∨ (*uw*)*x* ⊆ *D*]

Less formally, the last three properties respectively mean:

- if (*uv*)*w*, then not (*uw*)*v* and not (*wv*)*u*
- if (*uv*)*x* and (*vw*)*x*, then (*uw*)*x*
- if (*uv*)*x*, then either (*uv*)*w*, or (*uw*)*x*, or both

To conclude, suppose that *D* implies *k* triplets *t* _1_, …, *t*_*k*_ such that *k* ≥ 1.

In this case, one has

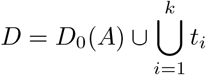

and either

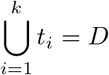

for the case where for all distinct *x, y* ∈ *A*, there are at least one triplet *t* ⊆ *D* and one *z* ∈ *A* such that *t* ∈ {(*xy*)*z*, (*xz*)*y*, (*yz*)*x* }, or

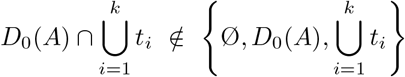

otherwise. As a corollary, we define *T* (*D*) as the set of all the triplets included into *D*. In other words,

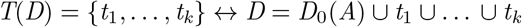

## 3 Consistency and ahograph-based representation of a triplet set

### 3.1 Consistency of a triplet set

First, let *l* (*t*) =_df_ {*x, y, z* } for any triplet *t* such that *t* ∈ {(*xy*)*z*, (*xz*)*y*, (*yz*)*x* }. Then, let *T* = {*t* _1_, …, *t*_*k*_} be a set of *k* triplets such that *k* ≥ 1. In this case, the set *L*(*T*) of all the objects related by the triplets of *T* can be defined as follows:

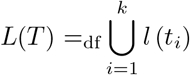

*T* is thus *consistent* if and only if the triplets of *T* are *k* to *k compatible*, i.e., if and only if there is at least one *D* ∈ **D**(*A*) such that *A* ⊇ *L*(*T*) and *T* (*D*) ⊇ *T*. Naturally, *T* is trivially consistent when *k* = 1. Given any two triplets *t* and *t*’,*t* is also *pairwise compatible* with *t*’ if and only if {*t, t*’} is consistent. Therefore, *T* is consistent only if *t* is pairwise compatible with *t*’ for all *t, t*’ ∈ *T* (note that the converse is false; for example, {(*ab*)*c*, (*bc*)*d*, (*cd*)*a*} is inconsistent, although {(*ab*)*c*, (*bc*)*d* }, {(*ab*)*c*, (*cd*)*a*}, and {(*bc*)*d*, (*cd*)*a*} are all consistent).

### 3.2 Undirected simple graph

A (*finite undirected simple*) *graph* is an ordered pair *G* = ⟨*V, E* ⟩ such that *V* is a finite non-empty set and *E* ⊆ {*X* ⊆ *V* | |*X* | = 2}. If appropriate, the elements of *V* and *E* are respectively called the *vertices* and the (*undirected*) *edges* of *G*. Then, a (*finite*) *edge-labeled (undirected simple) graph* is a 3-tuple *G* = ⟨*V, E, f* ⟩ for which ⟨*V, E* ⟩ is a graph, and *f* is a function from *E* to a non-empty set *L* of labels.

Given a graph *G* = ⟨*V, E* ⟩, two distinct vertices *x* and *y* of *G* are *connected* if and only if there is at least one *path* between *x* and *y*, i.e., if and only if there are *k* vertices *v* _1_, *v* _2_, …, *v*_*k*_ in *G* such that *v* _1_ = *x, v*_*k*_ = *y*, and {*v*_*i*_, *v*_*i*+1_} ∈ *E* for any *i* ∈ {1, …, *k* – 1}. *G* is then *connected* if and only if there is always at least one path between any two distinct *x, y* ∈ *V. G* is finally *complete* if and only if every pair of distinct vertices of *G* is connected by a single edge.

Let *G* = ⟨*V, E* ⟩ be a graph. An *induced subgraph* of *G* is any graph *H* = ⟨*W, F* ⟩ such that *W* ⊆ *V, F* ⊆ {{*x, y* } ∈ *E* | *x* ∈ *W* ∧*y*∈ *W* }, and for all *x, y* ∈ *W*, {*x, y* } ∈ *E* only if {*x, y* } ∈ *F*. From this, a *clique* of *G* (if it exists) is any complete subgraph of *G*.

### 3.3 The ahograph-based representation of a triplet set

Let *T* be a finite non-empty triplet set and let *X* ⊆ *L*(*T*) such that *X* ≠ ∅. The *ahograph* of *T* and *X* is then an edge-labeled graph AG(*T, X*) = ⟨*X, E*_*X*_, *f*_*X*_⟩ such that:

- for any *e* ∈ *E*_*X*_, there are exactly two distinct *x, y* ∈ *X* and at least one *z* ∈ *X* such that *e* = {*x, y* } and (*xy*)*z* ∈ *T* ; and
- *f* _X_ : *E*_*X*_ → {*Y* ⊆ *X* | 1 ≤ |*Y* | ≤ |*X*| − 2} is a function such that for any {*x, y* } ∈ *E*_*X*_ and any *z* ∈ *X, z* ∈ *f*_*X*_({*x, y* }) if and only if (*xy*)*z* ∈ *T*.

In other words, AG(*T, X*) is a graph such that (Bryant and Steel 1995: 439; Seemann and Helmuth 2018: 386): (i) *X* is its set of vertices; (ii) for all *x, y* ∈ *X*, there is exactly one undirected edge linking *x* and *y* if and only if there is at least one *z* ∈ *X* such that (*xy*)*z* ∈ *T* ; and (iii) any edge linking the vertices *x* and *y* is labelled by the set {*z* ∈ *X* | (*xy*)*z* ∈ *T* }.

Let AG(*T, X*) be an ahograph. A *component* of AG(*T, X*) is either a singleton set {*x* } ⊆ *X*, or any non-empty set *Y* ⊆ *X* such that |*Y* | *>* 1 and the induced subgraph ⟨*Y, E*_*Y*_, *f*_*Y*_⟩ of AG(*T, X*) is connected. For any finite non-empty triplet set *T*, it follows that *T* is consistent if and only if for any *X* ⊆ *L*(*T*) such that |*X* | ≥ 3, AG(*T, X*) is disconnected, i.e., it has at least two disjoint components (Aho et al. 1981; Bryant and Steel 1995: theorem 2; Seemann and Helmuth 2018: theorem 2.1).

## 4 Closure of a consistent triplet set

### 4.1 Closure of a set

For any non-empty set *U*, a *closure operator* on *U* is any function

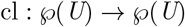

such that for all *X, Y* ⊆ *U* :

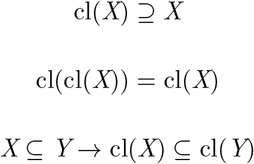

If appropriate, and for any *X* ⊆ *U*, cl(*X*) is the *closure* of *X*, and *X* is a *closed set* (relatively to cl) if and only if cl(*X*) = *X*.

### 4.2 Bryant and Steel’s definition

Let *T* be a finite non-empty triplet set. In this case, one can define the set co(*T*) of all the degree of equivalence *X* such that *T* (*X*) includes *T* as follows:

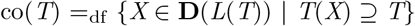

Thus, co(*T*) = ∅ if and only if *T* is inconsistent. Then, assume that *T* is consistent. In this case, the *closure* of *T* is the set cl(*T*) such that (Bryant and Steel 1995: 441; Grünewald *et al*. 2007: 523; Seemann and Helmuth 2018: 387):

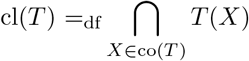

Naturally, that means that *T* is closed (for this closure operator) if and only if cl(*T*) = *T*. However, that also means that *T* is *combinable* if and only if there is exactly one *D* ∈ **D**(*L*(*T*)) such that *T* (*D*) = cl(*T*). Moreover, *T* is *complete* if and only if there is exactly one *D* ∈ **D**(*L*(*T*)) such that *T* (*D*) = *T*. As a result, *T* is complete only if *T* is both closed and combinable. Conversely, *T* is consistent if *T* is either closed, combinable, or both.

### 4.3 Redefinition in terms of ahograph

For all distinct *x, y, z* ∈ *L*(*T*), (*xy*)*z* ∈ cl(*T*) if and only if there is at least one *X* ⊆ *L*(*T*) such that AG(*T, X*) contains exactly two components *A* and *B* such that {*x, y* } ⊆ *A* and *z* ∈ *B* (Seemann and Helmuth 2018: theorem 2.6).

Let *T* be a finite non-empty consistent triplet set, and let (*xy*)*z* such that *x, y, z* ∈ *L*(*T*). In this case,

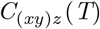

is the set of all pair {*X, Y* } for which *X, Y* ⊆ *L*(*T*) and AG(*T, X* ∪ *Y* } has exactly two components *X* and *Y* such that *x, y* ∈ *X* and *z* ∈ *Y* (Seemann and Helmuth 2018: Definition 3.2). Then assume that (*xy*)*z* ∈ *T*. In this case,

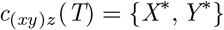

is the unique element of *C* _(*xy*)*z*_ (*T*) such that |*X* ^*^ ∪ *Y* ^*^| ≥ |*X* ∪ *Y* | for all {*X, Y* } ∈ *C* _(*xy*)*z*_ (*T*) (Seemann and Helmuth 2018: Definition 3.6). After that, let 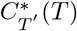 such that for any *T*’ ⊆ cl(*T*) (Seemann and Helmuth 2018: Definition 3.6):

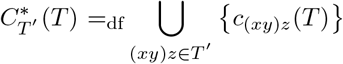

As a result, *T* is a non-empty consistent triplet set only if (Seemann and Helmuth 2018: Theorem 3.9):

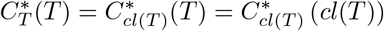

Finally, let

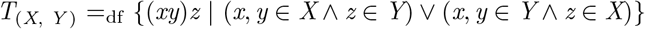

for any *X, Y* ⊆ *L*(*T*). In this case (Seemann and Helmuth 2018: Theorem 5.1):

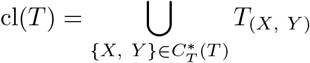

## 5 A finite set of rules to compute the closure

In this part, we propose to compute cl(*T*) only from the basic properties of triplets listed above. Our strategy is to deduce a 1-2 transitive closure and a 2-3 negatively transitive closure, and then to prove that a specific combination of them is equivalent to Bryant and Steel’s closure.

### 5.1 1-2 transitive closure

Let *T* be a consistent, finite, and non-empty triplet set. For all distinct *w, x, y, z* ∈ *L*(*T*), the 1-2 transitivity property means that:

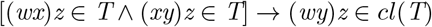

As a result, one can define the 1-2 transitive closure cl_T_(*T*) of *T* as follows:

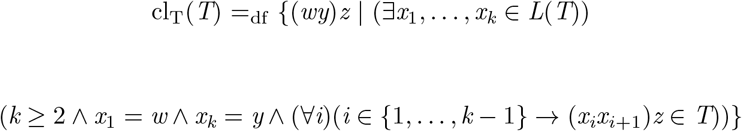

In terms of ahograph, the main consequence of this closure is that for any {*X, Y* } 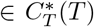 and all distinct *w, y, z* ∈ *L*(*T*) such that *w, y* ∈ *X* and *z* ∈ *Y*, it adds a *z* -labeled edge between *w* and *y* each time there is a *z* -labeled path from *w* to *y*.

### 5.2 2-3 negatively transitive closure

Now, let *T* be a consistent, finite, a nd non-empty triplet set. Under the assumption that *T* is true, it follows that there is necessarily exactly one degree of equivalence *D* on *L*(*T*) such that *T* (*D*) ⊇ *T*. For all distinct *w, x, y, z* ∈ *L*(*T*), the 2-3 negative transitivity property then means that:

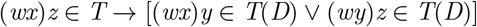

In addition, the pairwise incompatibility property means that for all *x, y, z* ∈ *L*(*T*):

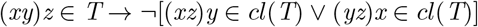

(Since *T* ⊆ cl(*T*) ⊆ *T* (*D*), it follows three possibilities:

- if (*wx*)*z* ∈ *T* and (*wx*)*y* ∈*/* cl(*T*), then (*wy*)*z* ∈ cl(*T*) and (*xy*)*z* ∈ cl(*T*)
- if (*wx*)*z* ∈ *T* and (*wy*)*z* ∈*/* cl(*T*), then (*wx*)*y* ∈ cl(*T*)
- if (*wx*)*z* ∈ *T* and (*xy*)*z* ∈*/* cl(*T*), then (*wx*)*y* ∈ cl(*T*)

Remarkably, each of these three inferences can be formulated in terms of ahograph. Indeed, the first one means that there is a cycle *w* -*x* -*y* -*w* with edges labeled by *z* whenever there is an edge labeled by *z* between *w* and *x* and no edge labeled by *z* between *x* and *y*. The second one then means that there is an edge labeled both by *y* and *z* whenever there is an edge labeled by *z* between *w* and *x* and no edge labeled by *z* between *w* and *y*. As for the last one, it leads to the same conclusion as the previous whenever there is an edge labeled by *z* between *w* and *x* and no edge labeled by *z* between *x* and *y*.

The 2-3 negatively transitive closure of *T* rests on the last two inferences. For this, let *Z* ⊆ *L*(*T*), and let *X, Y* ⊆ *Z*. One may then define the set *T*_*Z*/(*X, Y*)_ from AG(*T, Z*) as follows:

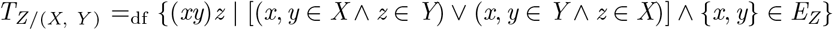

In this case, the negatively transitive closure of *T* is finally the set cl_NT_(*T*) such that:

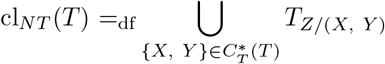

The main consequence of this closure is that it adds all the possible vertices of *Y* as labels of the edges of *X* and conversely for any {*X, Y* }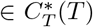.

### 5.3 Closure of a consistent triplet set

Let *T* be a finite, non-empty, and consistent triplet set. In this case,

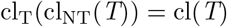

implies that the closure of a consistent triplet set can be reach by application on a set of the 2-3 negatively transitive closure, and then of the 1-2 transitive closure (Fig. 1).

**Fig. 1.**
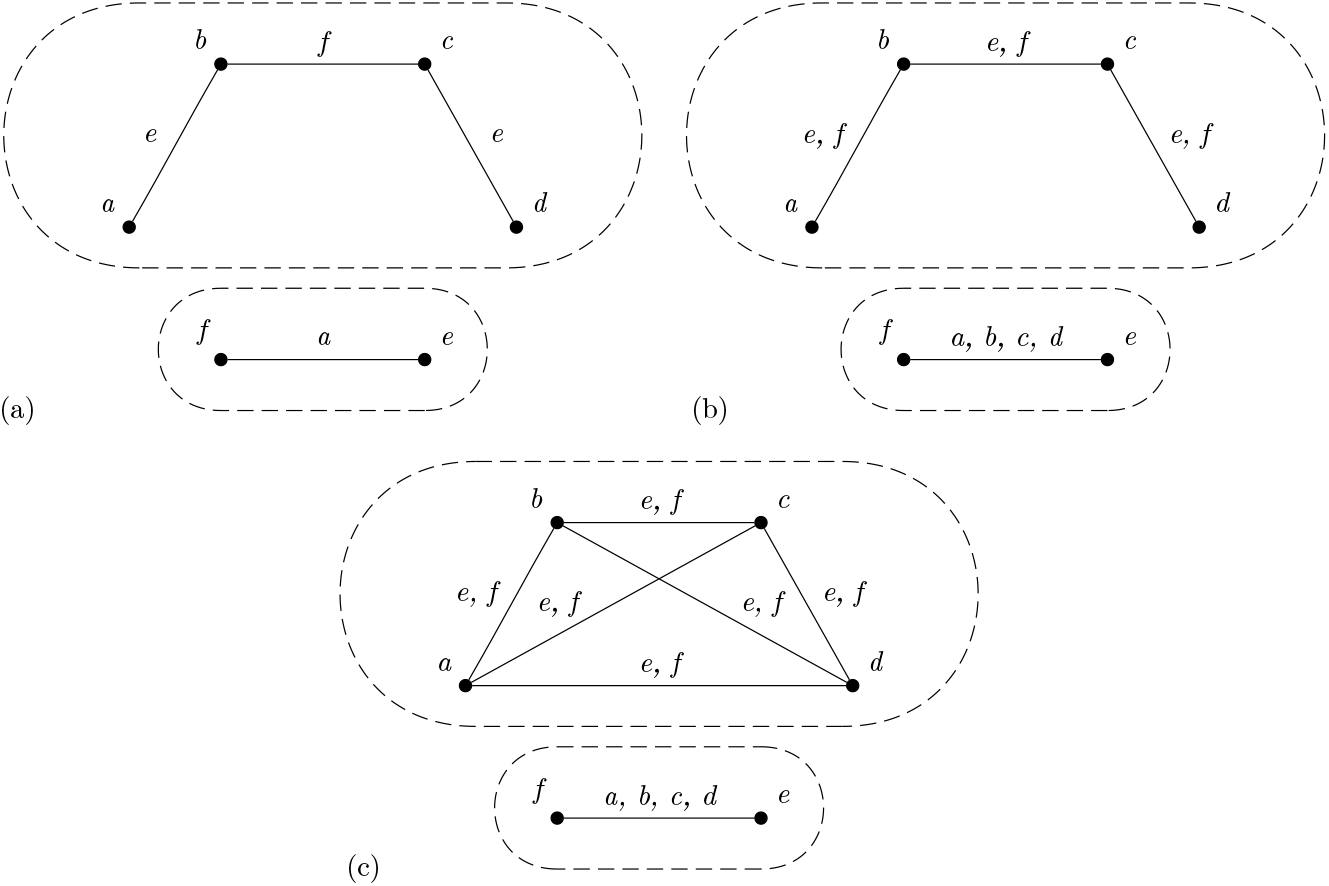
Representation of triplet sets by ahographs. (a) Representation of a 3is set *T* = {(*ab*)*e*, (*bc*)*f*, (*cd*)*e*, (*ef*)*a}*. (b) Representation of the negatively transitive closure cl_NT_(*T*) of *T*, with new labels on branches representing six additional triplets relatively to *T*. Each branch is labeled by all items from the opposite component. (c) Representation of the double closure cl_T_(cl_NT_(*T*)) of *T* with new branches and leading to a complete graph for each component where each branch is labeled with all items from the opposite component. The dotted lines indicate the two components *X* and *Y* of *AG*(*T, Z*).

**Proof**. We first begin with cl_T_(cl_NT_(*T*)) ⊆ cl(*T*). Let three distinct *x, y, z* ∈ *L*(*T*). Then, assume that (*xy*)*z* ∈ cl_T_(cl_NT_(*T*)). In this case, there is at least one ahograph AG(*T, Z*) such that *Z* ⊆ *L*(*T*) and with exactly two disjoint components *X, Y* ⊆ *Z* such that *x, y* ∈ *X* and *z* ∈ *Y*. But if true, then (*xy*)*z* ∈ cl(*T*).

We now turn with cl(*T*) ⊆ cl_T_(cl_NT_(*T*)). Let again three distinct *x, y, z* ∈ *L*(*T*) and assume (*xy*)*z* ∈ cl(*T*). In this case, there is at least one ahograph AG(*T, Z*) such that *Z* ⊆ *L*(*T*) and with exactly two disjoint components *X, Y* ⊆ *Z* such that *x, y* ∈ *X* and *z* ∈ *Y*. But if true, then there is necessarily one path between *x* and *y* into AG(*T, Z*). 2-3 negative transitivity then implies that each edge of the path connecting *x* and *y* in AG(cl_NT_(*T*), *Z*) is labelled by *Y*. In doing so, 1-2 transitivity finally implies that *x* and *y* are directly connected by a *Y* -labeled edge. As a result, (*xy*)*z* ∈ cl_T_(cl_NT_(*T*)).

## Acknowledgements

We thank Mark Wilkinson for sharing with us a stimulating example of five three-item statements, and Mike Steel for sending us Dekker’s remarkable dissertation. We are also grateful to Paul Zaharias for his helpful comments on the manuscript. Finally, we thank Rachel Vautrin, Rene Zaragueta, Paul Zaharias, and all the members of the Paris working group on 3ia and Cladistics for fruitfull discussions.

## References

Adams EN (1986) N-trees as nestings: Complexity, similarity, and consensus. Journal of Classification 3:299–317. 10.1007/BF01894192

Aho AV, Sagiv Y, Szymanski TG, Ullman JD (1981) Inferring a tree from lowest common ancestors with an application to the optimization of relational expressions. SIAM Journal on Computing 10:405–421

Bininda-Emonds ORP (2004) The evolution of supertrees. Trends in Ecology Evolution 19:315–322. 10.1016/j.tree.2004.03.015

Bryant D (2003) A classification of consensus methods for phylogenetics. DIMACS series in discrete mathematics and theoretical computer science 61:163–184

Bryant D, Steel M (1995) Extension Operations on Sets of Leaf-Labelled Trees. Advances in applied mathematics 16:425–453

Camin JH, Sokal RR (1965) A method for deducing branching sequences in phylogeny. Evolution 19:311–326

Colonius H, Schulze HH (1981) Tree structures for proximity data. British Journal of Mathematical and Statistical Psychology 34:167–180

Dekker MCH (1986) Reconstruction methods for derivation trees. Department of Mathematics and Computer Science, Vrije Universiteit, Amsterdam

Dobson AJ (1974) Unrooted Trees for Numerical Taxonomy. Journal of Applied Probability 11:32–42

Grünewald S, Steel M, Swenson MS (2007) Closure operations in phylogenetics. Mathematical Biosciences 208:521–537.

Hennig W (1966) Phylogenetic Systematics. University of Illinois Press

Hull DL (1970) Contemporary Systematic Philosophies. Annual Review of ecology and systematics 1:19–54

Islam M, Sarker K, Das T, et al (2020) STELAR: a statistically consistent coalescent-based species tree estimation method by maximizing triplet consistency. BMC Genomics 21:136. 10.1186/s12864-020-6519-y

McMorris FR, Powers RC (2003) The arrovian program from weak orders to hierarchical and tree-like relations. In: Janowitz MF, Lapointe FJ, McMorris FR, et al. (eds) Bioconsensus, American Mathematical Society. Providence, pp 37–45

Nelson G (1994) Homology and systematics. In: Hall BK (ed) Homology: The Hierarchical Basis of Comparative Biology, Academic Press. San Diego, pp 101–149

Nelson G (1970) Outline of a theory of comparative biology. Systematic Zoology 19:373–384

Nelson G (1973) Classification as an expression of phylogenetic relationships. Systematic Zoology 22:344–359

Nelson G (1972) Phylogenetic Relationship and Classification. Systematic Zoology 227–231

Nelson G, Ladiges PY (1991) Three-area statements: standard assumptions for biogeographic analysis. Systematic Biology 40:470–485

Nelson G, Platnick NI (1991) Three-Taxon Statements: A More Precise Use of Parsimony? Cladistics 7:351–366

Nguyen N, Mirarab S, Warnow T (2012) MRL and SuperFine+MRL: new supertree methods

Prin S (2016) The relational view of phylogenetic hypotheses and what it tells us on the phylogeny/classification relation problem. In: Williams D, Schmitt M, Wheeler Q (eds) The Futur of Phylogenetic Systematics: The Legacy of Willi Hennig, Cambridge University Press. pp 431–468

Seemann CR, Hellmuth M (2018) The matroid structure of representative triple sets and triple-closure computation. European Journal of Combinatorics 70:384–407. 10.1016/j.ejc.2018.02.013

Sevillya G, Frenkel Z, Snir S (2016) Triplet MaxCut: a new toolkit for rooted supertree. Methods in Ecology and Evolution 7:1359–1365.

Vach W (1994) Preserving consensus hierarchies. Journal of Classification 11:59–77. 10.1007/BF01201023

Wilkinson M (1994) Common Cladistic Information and its Consensus Representation: Reduced Adams and Reduced Cladistic Consensus Trees and Profiles. Systematic Biology 43:343–368. 10.1093/sysbio/43.3.343

Wilkinson M, Cotton J, Thorley J (2004) The Information Content of Trees and Their Matrix Representations. Systematic Biology 53:989–1001. 10.1080/10635150490522737

